# Digital Manufacturing of Functional Ready-to-Use Microfluidic Systems

**DOI:** 10.1101/2023.05.08.539659

**Authors:** Vahid Karamzadeh, Ahmad Sohrabi-Kashani, Molly Shen, David Juncker

## Abstract

Digital manufacturing (DM) strives for the seamless manufacture of a functional device from a digital file. DM holds great potential for microfluidics, but requirements for embedded conduits and high resolution beyond the capability of common manufacturing equipment, and microfluidic systems’ dependence on peripherals (e.g. connections, power supply, computer), have limited its adoption. Microfluidic capillaric circuits (CCs) are structurally-encoded, self-contained microfluidic systems that operate and self-fill thanks to precisely tailored hydrophilicity. CCs were heretofore hydrophilized in a plasma chamber, but which only produces transient hydrophilicity, lacks reproducibility, and limits CC design to open surface channels sealed with a tape. Here we introduce the additive DM of monolithic, fully functional and intrinsically hydrophilic CCs. CCs were 3D printed with commonly available light engine-based 3D printers using polyethylene(glycol)diacrylate-based ink co-polymerized with hydrophilic acrylic acid crosslinkers and optimized for hydrophilicity and printability. A new, robust capillary valve design and embedded conduits with circular cross-sections that prevent bubble trapping are presented, and complex interwoven circuit architectures created, and their use illustrated with an immunoassay. Finally, the need for external paper capillary pumps is eliminated by directly embedding the capillary pump in the chip as a porous gyroid structure, realizing fully functional, monolithic CCs. Thence, a computer-aided design file can be made into a CC by commonly available 3D printers in less than 30 minutes enabling low-cost, distributed, DM of fully functional ready-to-use microfluidic systems.

## 1. Introduction

Digital manufacturing (DM) implies the largely automated, computer-centric fabrication of customizable products from a digital file to a final product. Widespread adoption of DM requires both widely available manufacturing equipment, and designs that can be directly manufactured by these equipment. Microfluidic systems have traditionally been made by clean room microfabrication and replication methods^[1]^, but the adoption of classical and emerging subtractive and additive manufacturing processes for microfluidics fabrication makes it a candidate for DM.^[2]^ Subtractive processes include CNC milling^[3]^ and laser cutting^[4]^ while additive processes include 3D printing using filament-based printing^[5,6]^ and light-based photopolymerization^[7–9]^ have become popular over the last few years owing to rapid technological progress, low capital costs, and low skill requirements^[10]^. In particular, application-specific microfluidic chips would benefit from on-demand and on-the-fly design and manufacturing. DM could drive a broader adoption of microfluidics if one could print functional microfluidic systems similar to the way one can print out a digital text and art, such as this manuscript, on one’s own printer at the office or home in a matter of minutes.

There are several challenges that prevent the rapid adoption of DM for microfluidics. Firstly, most microfluidic systems are computer controlled, and require peripherals that can be costly and bulky, and are application specific, and are thus not universally available. This requirement doubly impacts the adoption of DM because it is conditional on users pre-owning peripheral control systems and will limit the use of DM to making microfluidic chips. Moreover, the very use of computers and peripherals means that what is commonly referred to as active microfluidics^[11]^ is in actuality based on generic, passive chips that can be used for a variety of microfluidic applications by reprogramming the computer without the need for physically reconfiguring the chip itself. As mass production of generic chips is cheaper than DM, it remains advantageous relative to DM. Secondly, many microfluidic systems cannot be made via DM because DM-compatible manufacturing processes lack versatility. Indeed, if the chip architecture is very complex, or the chips require special materials, or different materials, or if they require microscopic features, or a combination of both microscopic and macroscopic features, or special surface chemical properties, then they may not be compatible with current DM processes.

Capillary microfluidics can operate without requiring any bulky peripherals by making use of capillary phenomena defined by the geometry and surface chemistry of microchannels to control fluid flow^[12,13]^. Additive manufacturing of monolithic, functional capillary microfluidics using a porous material with different powders of variable hydrophilicity to create microfluidic ‘channels’ was recently demonstrated using a customized binder jet 3D printer^[14]^. However, powder printing and this design were dependent on a custom printer design, thus preventing broader adoption. Stereolithography and digital light processor (DLP) have become widely available, and over the last few years, multiple groups^[15,16]^ have developed systems and processes for making microfluidic devices with high resolution and advanced functionality. Polyethylene(glycol)diacrylate (PEGDA) with a molecular weight of 250 (PEGDA-250) has emerged as a popular material for additive 3D printing of microfluidics devices^[17,18]^. Whereas the initial use of PEGDA was making ‘active’ microfluidics that uses passive chips dependent on peripherals,^[17,18]^ more recently, DLP printing has been used to make so-called capillaric circuits (CCs)^[12,19]^.

CCs are capillary microfluidics assembled from individual capillaric elements that structurally encode liquid handling algorithms. An initial study using replica molded CCs from DLP printed molds established the feasibility of DLP printing despite a lower resolution than clean room fabricated CCs^[20]^, and alleviated initial concerns on whether capillary flow functions such as filling and flow stop could be realized.^[20]^ More recently, application-specific CCs were directly printed on-demand and used to execute structurally pre-programmed liquid handling algorithms with as many as several hundred operations, thanks to the microfluidic chain reaction (MCR), and for applications in various assays.^[12,19,21]^ CCs operate thanks to precisely controlled hydrophilicity with a water contact angle between ∼30-60º to meet the dual requirements of self-filling by capillary flow, and flow stoppage at capillary stop valves (SVs)^[21]^. Currently, 3D-printed CCs require plasma post-processing for achieving the required hydrophilicity, which necessitates a plasma chamber, an equipment that is not readily available^[12]^. Moreover, this post-processing entails two additional limitations: hydrophilicity is fleeting and prevents CCs storage and field usage, and only open, surface channels can be plasma-activated, and hence CCs are printed as open surface structures enclosed with a hydrophobic cover – hydrophobicity is indeed required to preserve SVs function. The cover also needs to be cut-out for sample delivery. In short, there remained multiple impediments that prevented direct DM of CCs.

Here, we introduce a modified PEGDA-250 ink that is intrinsically hydrophilic for DM of monolithic, fully functional CCs with embedded conduits. We further introduce new designs for SVs and non-rectangular cross-sections that both improve the functionality and reliability of 3D-printed CCs. CCs with embedded interwoven conduits were used for serial delivery of reagents using the MCR and capillary retention burst valves (RBVs)^[22]^, and validated their use in immunoassays using SARS-CoV-2 protein. Finally, capillary pumps made from high-resolution, mathematically-generated geometries were directly integrated into the CCs, circumventing the need for external paper pumps. Thanks to these advances, fully functional CCs were printed within less than 30 minutes on a commercial DLP 3D printer, paving the way for distributed DM of functional microfluidic systems.

## 2. Results and Discussions

**Figure 1** illustrates the streamlined, rapid DM of monolithic CCs with an intrinsically hydrophilic, photopatternable ink (CCInk) comprising four main constituents, namely a monomer (PEGDA-250), a photoinitiator (diphenyl(2,4,6-trimethylbenzoyl) phosphine oxide, TPO), a photoabsorber (isopropylthioxanthone, ITX), and a hydrophilic agent and cross-linker (acrylic acid, AA).

**Figure 1.**
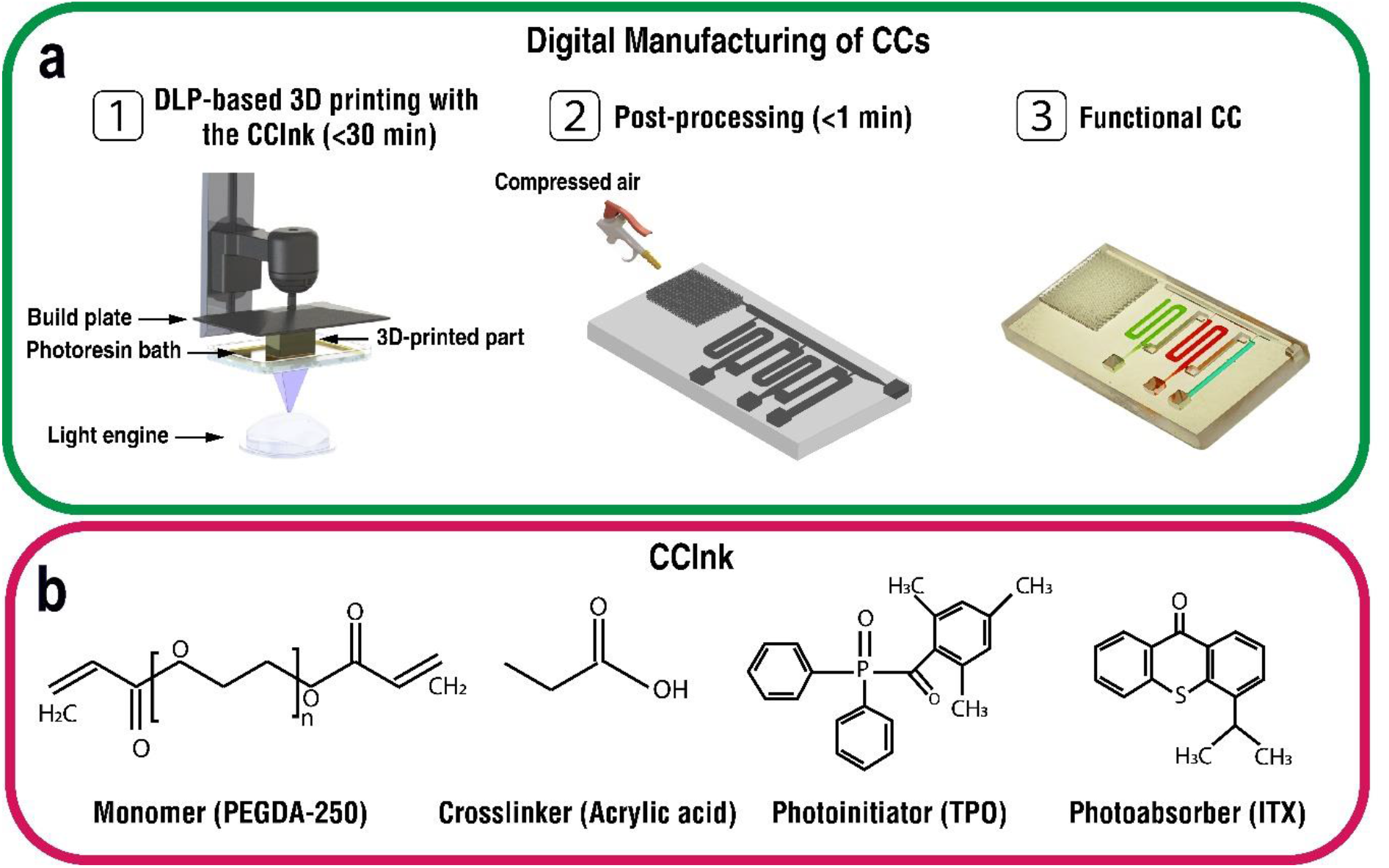
Digital manufacturing (DM) of functional capillaric circuits (CCs) using an intrinsically hydrophilic ink (CCInk). **(a)** 1) Additive manufacturing of CCs from a digital design file using a light engine and layer-by-layer vat photopolymerization. 2) Removal of uncured CCInk under a stream of compressed air (<1 min). 3) Functional CC loaded with reagents and primed to execute the structurally encoded (*i*.*e*. pre-programmed) capillary flow events following the addition of the triggering solution. **(b)** CCInk composition including polyethyleneglycol diacrylate (PEGDA-250) monomer, acrylic acid (AA) additive to tune hydrophilicity, a photinitiator (TPO) and a photoadsorber (ITX).

### 2.1. Formulation and characterization of CCInk

CCs made of native PEGDA-250 have a contact angle of ∼65°, which albeit considered hydrophilic, is too hydrophobic for reliable self-filling of microchannels by aqueous solutions, especially in the presence of hydrophobic cover. While it is also possible to plasma-treat PEGDA-250 and increase hydrophilicity, the surface gradually returns to its original low-hydrophilicity state over the course of 4h (**Figure S1, Supporting Information)**.

To permanently increase the hydrophilicity of CCs towards supporting reliable self-filling, we incorporated acrylic acid (AA) and methacrylic acid (MA) into the PEGDA ink^[23,24]^. AA and MA both have a vinyl group and can thus be co-polymerized with PEGDA by photopolymerization. The acidic groups of AA and MA can form hydrogen bonds with water, and thus lowering the contact angle and increasing hydrophilicity.^[23,24]^ We reasoned that by adjusting the ratio of PEGDA:Additive, the hydrophilicity could be adjusted to fall within the optimal range for autonomous, structurally encoded flow operations by CCs.

The contact angle of water on 3D-printed parts as a function of the percentage of AA and MA was measured. **Figure 2a** shows that it can be lowered to ∼35° for 10% or higher additive concentrations. We tested higher concentrations of additives, but we observed no further changes in the contact angle while printing resolution was compromised as additive-additive cross-linking competes with additive-PEGDA cross-linking (results not shown). Toluidine blue is an acidophilic dye that reversibly binds to acidic molecules in a pH-dependent manner and was previously used to characterize carboxyl groups. We utilized toluidine blue assay (TBO) to quantify the density of carboxyl groups in 3D-printed structures (**Figure 2b)**. The density of COOH as a function of AA and AM concentration in the ink shows saturation once the concentration reaches 10%. Based on these results, CCInk was formulated with 10% of AA because a contact angle of 35° meets the opposing requirement of self-filling by capillary flow and of halting it at SVs. To evaluate the potential for storage of CCs made with CCInk, we assessed the long-term stability of CCInk by measuring the contact angle with water over time on 3D-printed parts stored at room temperature over a period of 16 weeks, **Figure 2c**. We found the contact angle to be stable throughout, thus confirming the possibility of printing CCs, and using them at a later time.

**Figure 2.**
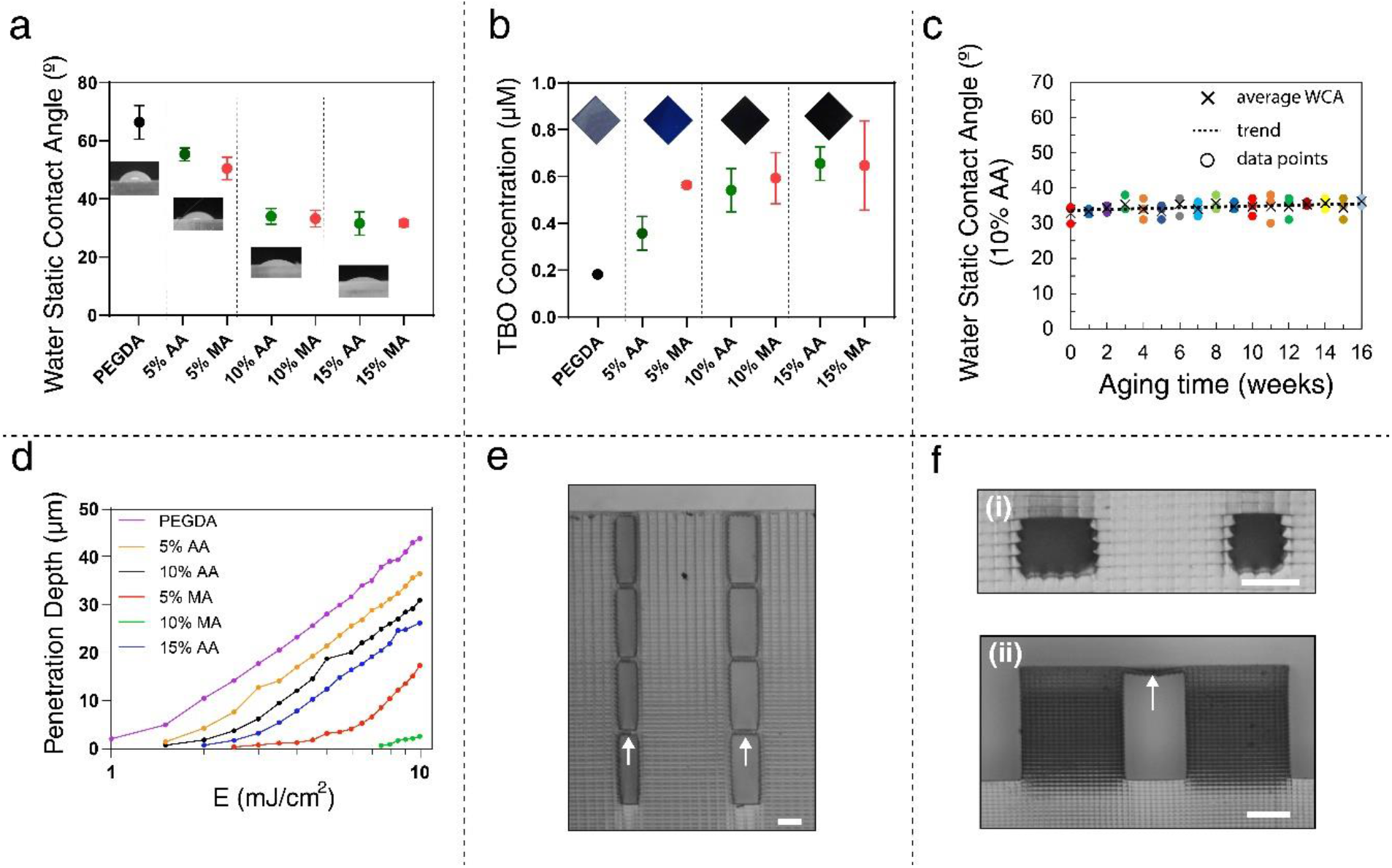
Photocurable ink design, optimization and 3D printability. **(a)** Static water contact angle for PEGDA-250 with different AA and MA concentrations. **(b)** Toluidine blue assay (TBO) quantification of carboxyl group density on 3D-printed samples. (**c)** serial contact angle measurements over a period of >16 weeks illustrating surface stability (the dashed line shows the linear trend of the average WCA). **(d)** Penetration depth characterization of the hydrophilic formulation with different co-monomer concentrations. **(e)** 3D-printed stacked microchannels separated with 25 μm membranes (white arrow) using a formulation with 10% AA. **(f)** (i) Embedded rectangular channels with dimensions of 100 ×135 μm ^2^ on the left and 100 × 108 μm ^2^ on the right, and (ii) 3D printed membrane with single layer roof membrane (indicated by the white arrow) using a formulation with 10% AA - Scale bar: 100 μm.

The photoadsorber and ink exhibit high efficiency at a wavelength of 385 nm, which matches the UV spectrum of 385 nm DLP light engines (**Figure S2a, Supporting Information)**. The ink’s absorbance remains unaffected by variations in AA and MA concentrations (**Figure S2b, Supporting Information)**. The manufacture of embedded conduits and thin bottom and ceiling walls depends on controlling the cross-linking depth and adsorption of UV light. The cross-linking depth was determined by applying uncured CCInk to a glass slide and exposing it to various light intensities shined through the glass. Upon removal of uncured ink, the thickness of cured ink can be measured, **Figure 2d**. As expected, an increase in additive concentration leads to a reduced cross-linking depth. Furthermore, according to the penetration depth results, PEGDA-AA required a lower exposure time for solidification than PEGDA-AM due to more effective cross-linking between AA and PEGDA. The reduced cross-linking in formulations with higher concentrations of AA and MA could be attributed to the lower molarity of acrylate groups, as the absorbance of the ink remains unaffected by variations in AA and MA concentrations. Hence, CCInk was formulated with PEGDA-AA and is the one used in the subsequent sections.

While the resolution of an open channel mainly depends on the projected pixel size in X and Y, embedded channels must also consider light penetration across the cross-linked membrane closing the channel and inadvertent crosslinking of entrapped ink immediately above it in Z. The ability to 3D print small embedded channels is crucial for the fabrication of functional CCs. In addition, we demonstrated the printing of stacked embedded microchannels with 25 μm roof membranes, using a formulation with 10% AA (**Figure 2e**). Our ability to 3D print small embedded circular and rectangular channels would enable the seamless fabrication of CCs.

### 2.2. Embedded Stop Valve Optimization

In order to develop functional fully 3D-printed CCs, the functionality and reliability of trigger valves (TVs) and SVs that are critical to confine liquids to conduits and reservoirs in the circuits need to be reconsidered^[21]^. Previous CCs were printed as open channels at the surface of the chip and were sealed with a hydrophobic cover forming the ceiling. In this scenario, the 3 hydrophilic side walls of the conduit ensured self-filling by capillary force, while the hydrophobic ceiling was needed for stoppage of the incoming liquid at SVs and TVs. In contrast, when the conduits are embedded within the CCInK, the channel ceiling in the closed channel is hydrophilic as well, thus jeopardizing the functionality of the valves, and of the entire circuit. Consequently, it was necessary to characterize and optimize SV designs before developing complex CCs. In the conventional SV/TV, the flow within conduits experiences abrupt geometrical changes at the capillary valve only on three sides (bottom, left, and right), and the leakage from the top is controlled by having a fairly hydrophobic top surface (sealing cover). Hence, the meniscus is pinned at the SV outlet edge. Thanks to embedded channels, in the face-centric TV/SV design, the flow could be pinned at valves by having a sudden change on all sides (including the top surface) by designing valves at the middle of the second (orthogonal) microchannel (main channel). In this fashion, the flow will stop at the SV/TV even if the ceiling is hydrophilic by controlling the expansion angle from the top, which is not attainable in conventional designs. In this design, the vertical gap plays a significant role in the valve performance, so having a sufficient vertical gap is necessary to hold liquids in reservoirs robustly for a longer time^[20]^. **Figure 3b** compares face-centric and conventional SV designs 3D-printed with CCInk. In the case of conventional design, the valve bursts from the top side immediately after loading, and the liquid started spilling to the main channel. However, the face-centric valves could hold the liquid for a long time without leakage. The possibility of 3D printing embedded capillaric valves allows for controlling the expansion angle (90° or even more) on each side of valves and creating stronger and more robust TVs/SVs.

3D printing enables the construction of vertical channels that are not possible or challenging to fabricate using conventional techniques, allowing to development of vertical SVs/TVs from top or bottom. **Figure 3b** further shows a vertical SV connecting the reservoir to the main channel from the top. The vertical valve held liquid in the reservoir for 20 min without leakage and was insensitive to gravity effects. The functionality of the conventional and face-centric SVs was also verified by studying capillary filling within functional connections using the level-set method and COMSOL simulation. As shown in **Figure 3a**, the conventional valve failed to stop the flow right after capillary-filling, and the flow spread to the top surface of the expansion chamber, while the embedded valve stopped the flow without leakage under the same boundary conditions.

**Figure 3.**
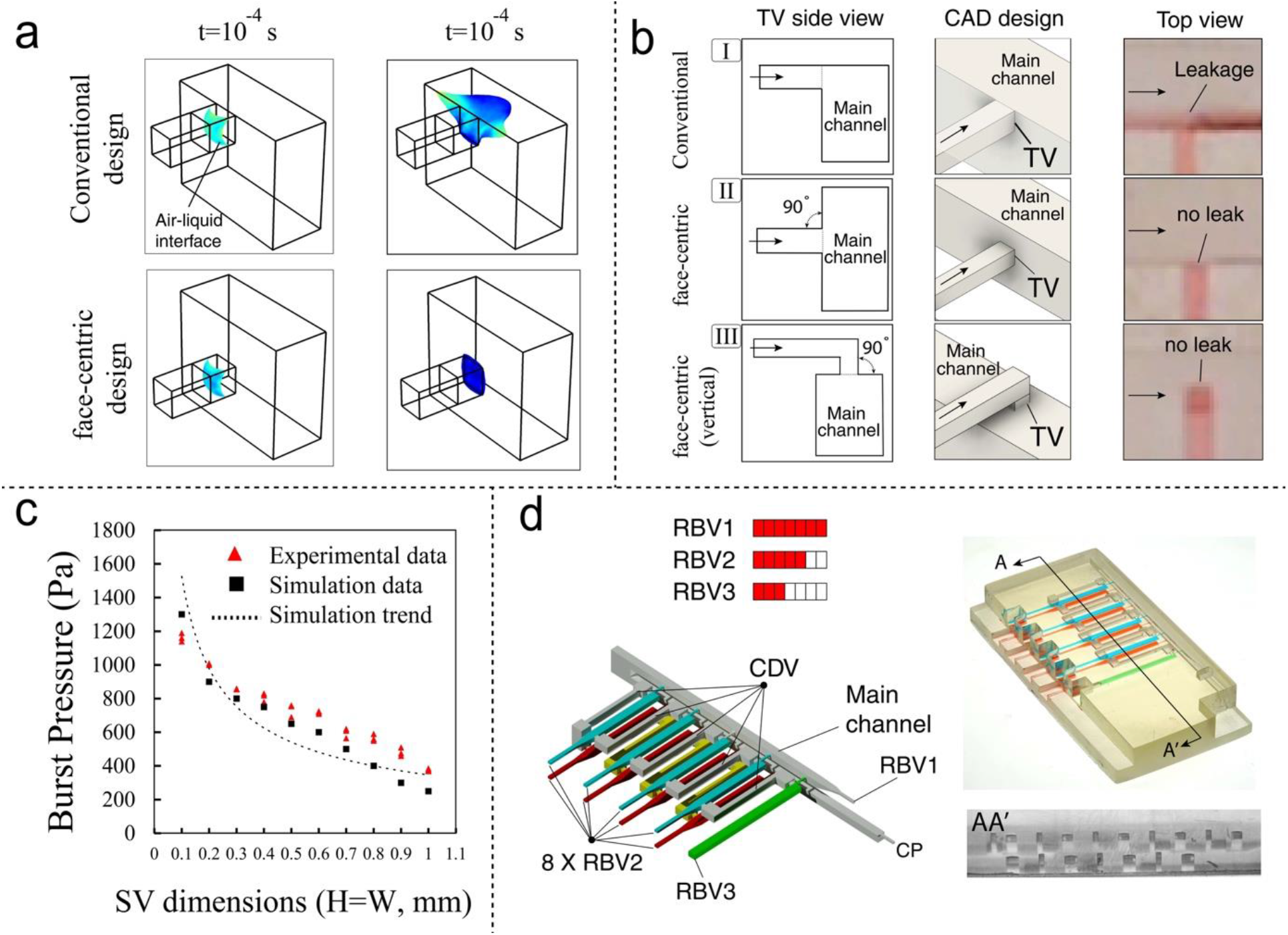
Redesigned and improved capillary stop valves. **(a)** FEM-based simulation of capillary flow reaching a conventional SV with the continuous ceiling (0*°* angle and 3 surfaces with 90*°* angle relative to flow direction) and of a face-centric SV (four 90*°* angles). For a CCInk with a water contact angle of40°, conventional SV/TV fail immediately while face-centric ones stop the liquid. **(b)** Experimental characterization of (I) conventional, and face-centric SV/TV intersecting the other conduit (II) laterally or (III) from the top; only face centric SV/TV could stop the liquid flow (arrows show the flow direction). **(c)** Comparison between simulation and experimental burst pressure for face-centric SV/TV with different square cross-section areas (from 100 to 1000 μm side) versus fitted values derived from a numerical model. **(d)** 3D printed two-layered CCs with face-centric elements for sequentially draining of 9 reservoirs using MCR (**See Video S1, Supporting Information**).

To further characterize the performance of the face-centric SVs, a pressure pump was used to measure bursting pressure at SVs of different sizes. Embedded valves with conduits with different square cross-section areas (from 100 to 1000 μm) were designed and 3D-printed for the burst pressure test, as shown in **Figure 3c**. Each SV was connected to a positive pressure line at the inlet and another line at the outlet to provide an adjustable pressure difference between the upstream and downstream of each SV. The pressure difference was gradually increased by applying a positive inlet pressure until the meniscus bursts on the sides of the SV. As shown in **Figure 3c**, the bursting pressure value decreased from ∼1200 Pa for the smallest SV to ∼350 Pa for the largest tested SV with a size of 1 mm. This suggests that face-centric SVs can withstand a higher pressure than conventional SVs (with a hydrophobic top surface) that were tested in our previous study^[12]^. Using finite element modelling, the bursting pressure at face-centric SVs with different cross-section areas was simulated and the highest pressure that could be rested by the SV calculated, finding a good concordance between experiments and theory.

As a proof-of-concept, a monolithic CC with embedded conduits, a two-layered CC with a MCR with nine capillary flow events, five on top and four on the bottom layers was manufactured (**Figure 3d, Video S1, Supporting Information)**. The 5^th^ reservoir in the top fluidic network is connected to the 6^th^ reservoir in the bottom network through a capillary domino valve (CDV), while the main conduit also bridges the two layers. Upon drainage of the reservoirs in the top layer, the ones in the bottom layer were drained. Manufacturing embedded channels could help integrate more conduits within the working footprint of the 3D printer. Vertical connections could add to the pressure head due to gravity, and interfere with capillary valve functionality, however as the height was low, functional CCs could be readily made.

### 2.3. 3D printing of CCs with embedded conduits with circular cross-sections

Microfluidic conduits with square and rectangular cross-sections are commonly used as they are easy to manufacture using standard microfabrication methods and 3D printing. However, for liquids where the contact angle falls < 45°, edge flow in the corners arises. Hence liquid filaments can reach an exit and clog it, thus trapping bubbles. This situation can arise in CCs when using low surface tension solutions, such as buffers including surfactants that are used in assays to prevent non-specific binding. When using a solution containing 0.1 % surfactant, with a contact angle of 25° with photocured CCInk, we observed corner flow and bubble trapping, **Figure 4a**. Therefore, we explored the possibility of printing conduits with circular cross-sections using vat polymerization 3D printing while considering the limited resolution of the 3D printer. To improve circularity and minimize artifacts, we utilized anti-aliasing techniques and a low layer thickness of 20 μm. Following optimization, we were able to successfully 3D print circular channels with diameters as small as 160 μm (**Figure 4b**), representing a slight trade-off in diameter compared to rectangular conduits, which were 108×100 μm^2^ (see **Figure 2f**). A serpentine conduit with circular cross-section with varying diameters was also filled with a low surface tension solution, and successfully averted bubble trapping, **Figure 4c**. A functional CC with three reservoirs with circular cross-sections each connected to RBVs^[21,22]^ with a different threshold was manufactured, **Figure 4d**. Each of the reservoirs was connected to the main conduit via a 200 μm diameter conduit, and upon filling and actuation, the three reservoirs were drained according to the pre-programmed sequence encoded by the RBV thresholds (**Videos S2-S3, Supporting Information**).

**Figure 4.**
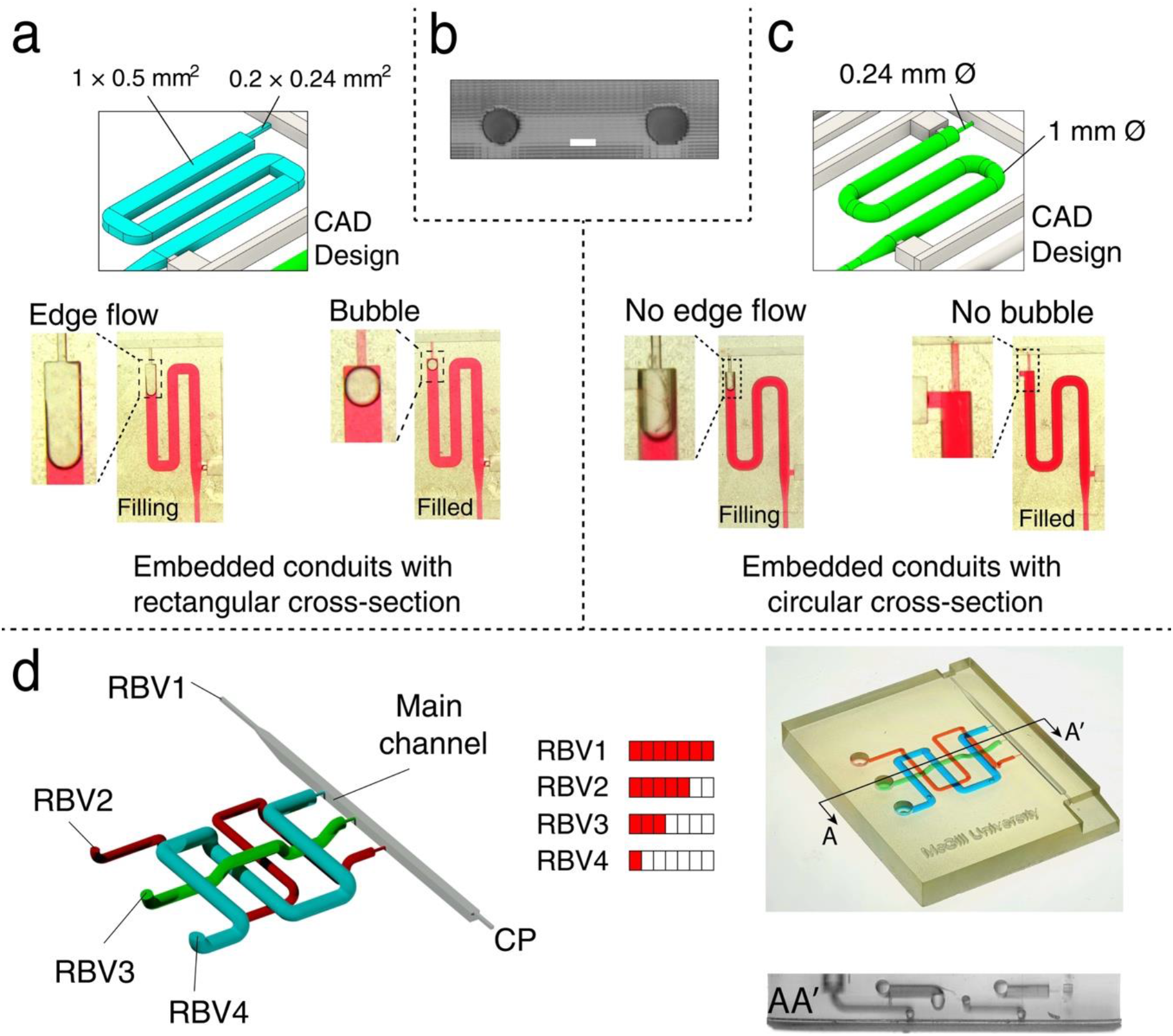
Monolithic 3D printed CCs with embedded conduits with circular cross-sections. **(a)** Flow of low surface tension liquid (0.1% Tween 20 in Mili-Q water) in a conduit with a rectangular cross-section is accompanied by edge flow and air bubble trapping. **(b)** Cross-sections of embedded circular conduits are shown: one with a diameter of 160 μm on the left, and one with a diameter of 200 μm on the right-scale bar is 100 μm **(c)** The same liquid in a circular conduit without edge flow and without bubble trapping. **(d)** 3D printed CC with circular cross-sections and face-centric SVs/TVs filled with dyed solutions in 3 reservoirs with 4 RBVs with different bursting thresholds for sequential delivery (**See Videos S2-S3**, Supporting Information).

### 2.4. Immunoassay with a 3D-Printed CC with embedded conduits

To demonstrate the power and compatibility of our 3D printing technique, a proof-of-concept ELISA assay was developed to detect SARS-CoV-2 antibodies. ELISAs require a sequence of steps to amplify the readout signal enzymatically. The designed ELISA protocol for the colorimetric detection of COVID-19 antibodies consists of three steps. First, the biotinylated antibody (sample) flows over the test zone where we already deposited the Nucleocapsid protein. Second, poly-HRP streptavidin flows over the test zone. Finally, the substrate flows over the test zone to chemically react with poly-HRP and generate a brown precipice visible to the naked eye (**Figure 5a**). We performed the ELISA assay manually (off-chip) to optimize the volume and concentration of reagents.

After optimizing capillaric elements and the assay, we designed and 3D-printed a monolithic CC microfluidic device to translate the manual ELISA assay into an autonomous multistep assay, in which sample and reagents are controlled and released sequentially to follow the assay protocol. This design includes three 3D reservoirs, one on the top layer and two on the bottom layer, to deliver the sample and reagents sequentially to the end of the main channel. The device is pre-programmed by designing 3D RBVs upstream of each reservoir, and all three are connected to the main channel through embedded functional connections (face-centric TVs) at their downstream to release the assay components sequentially. CCInk and 3D printing enable the manufacturing of compact devices by designing freeform capillaric elements and channels in three dimensions. Taking advantage of this characteristic, two layers of channels were designed and fabricated in this particular design, as shown in **Figure 5a**. A lateral flow module is connected to the end of the main channel. The module comprises a nitrocellulose membrane (test zone) sandwiched between two capillary pumps to ensure the flow throughout the assay. Before running the chip, the functionality of capillaric valves and sequential delivery were verified using dyed solutions, as shown in **Figure 5b**. All three reservoirs are first filled, and solutions are stopped into the functional connections. Then the assay starts by adding buffer into the main channel. The flow in each reservoir resumes when the functional connections are triggered by the trigger buffer in the main channel. Upon contact with the trigger buffer, solutions in reservoirs are released into the main channel and wicked sequentially through the main channel. Error! Reference source not found.**b** shows the sequential steps when the main channel is filled and connected to a paper pump (**Video S4, Supporting Information**). The blue solution (sample) in the top layer is released first, followed by releasing green and red solutions representing poly-HRP streptavidin and substrate, respectively, in the bottom layer. The CAD design of the fluidic network shows how channels were connected in 3D. Using a series of chips and samples at six different concentrations (0 to 10000 ng/ml) with three replicates of COVID-19 antibody in buffer, we established a binding curve. The assay results appeared as a line, which was scanned to quantify the signal, **Figure 5c**.

**Figure 5.**
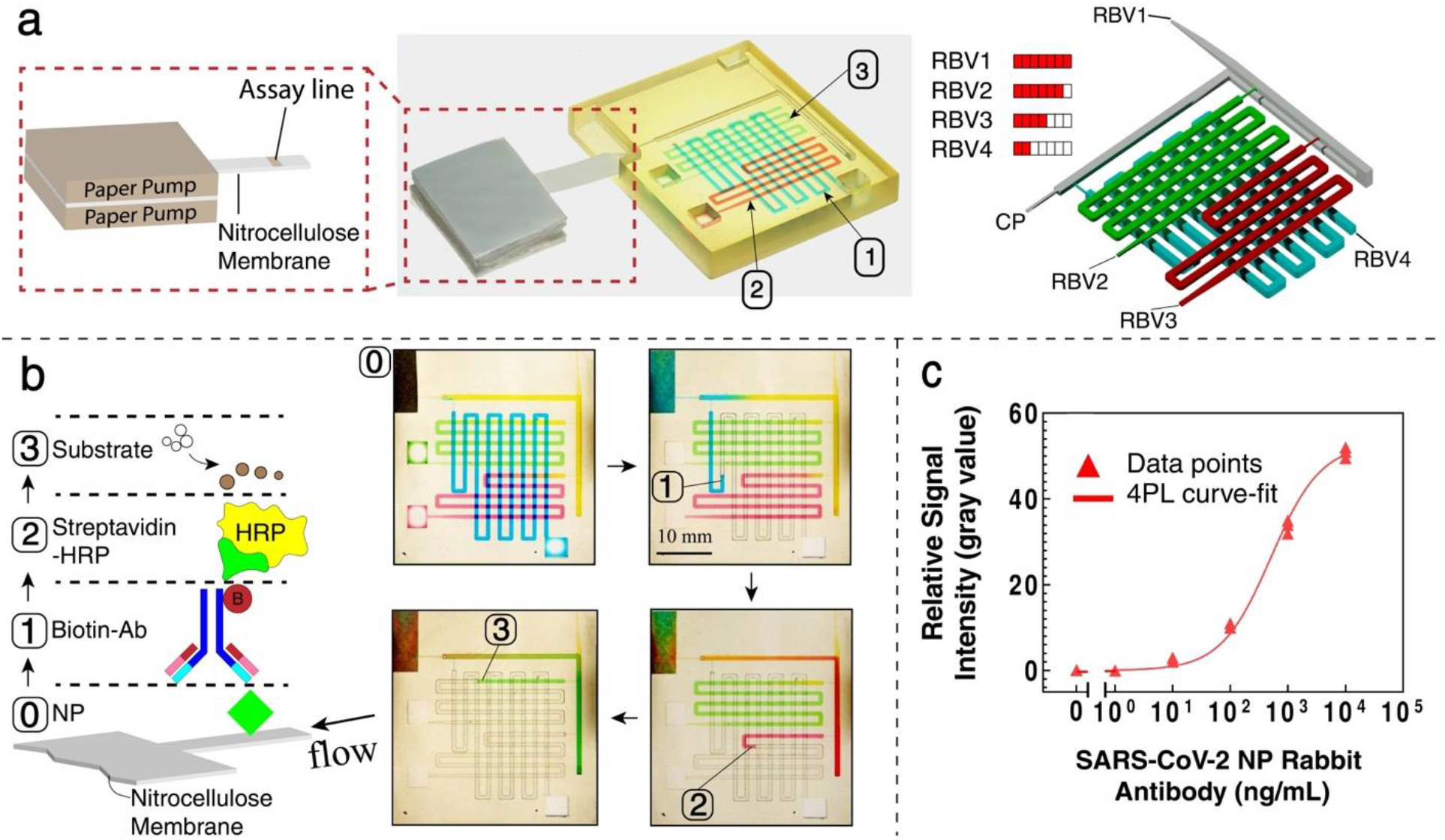
An autonomous assay using a 3D-printed monolithic CC. **(a)** A 3D-printed chip to deliver samples and reagents sequentially to the test zone where nucleocapsid protein (NP) is already spotted on the nitrocellulose membrane (NM) to complete the assay workflow. The assay includes three steps; biotinylated rabbit-anti NP (150 μl), Streptavidin-HRP (40 μl), and substrate (DAB, 50 μl) to produce colorimetric signals. **(b)** Sequential and pre-programmed release of colored solutions triggered by connecting the paper pump (**Video S4, Supporting Information**). **(c)** Assay results and the binding curve obtained by imaging the test zone with a scanner (fitted with a 4-paramater regression, 3 replicates for each point).

### 2.5. Full Additive Manufacturing of CC with Gyroid Capillaric Pump (GCP)

Using 3D Printing and CCInK, we manufactured fully functional circuits without the need for a paper pump. We adopted a gyroid^[25–27]^ based on triply periodic minimal surfaces (TPMS) as the capillary pump (GCP) (**Figure 6a**). The comparative high-resolution of the DLP printers enabled us to make embedded small features and form a capillary pump embedded integrated into the CC. The capillary pump is an essential functional unit of CCs that needs to meet opposing requirements, namely providing a sufficiently high capillary pressure to suck the liquid into the pump and provide a sufficiently large reservoir for liquids (serving also as the waste), while also not significantly adding to the flow resistance of the circuits. These requirements were thus often met by paper pumps, but which need to be clamped onto the CC. 2D capillary pumps have been made previously, notably by microfabrication, but the volumetric capacity was limited by the 2D geometry. 3D capillary pumps have not been widely explored, in part because the small feature size needed for high capillary pressure and the high open ratio were difficult to achieve.

**Figure 6.**
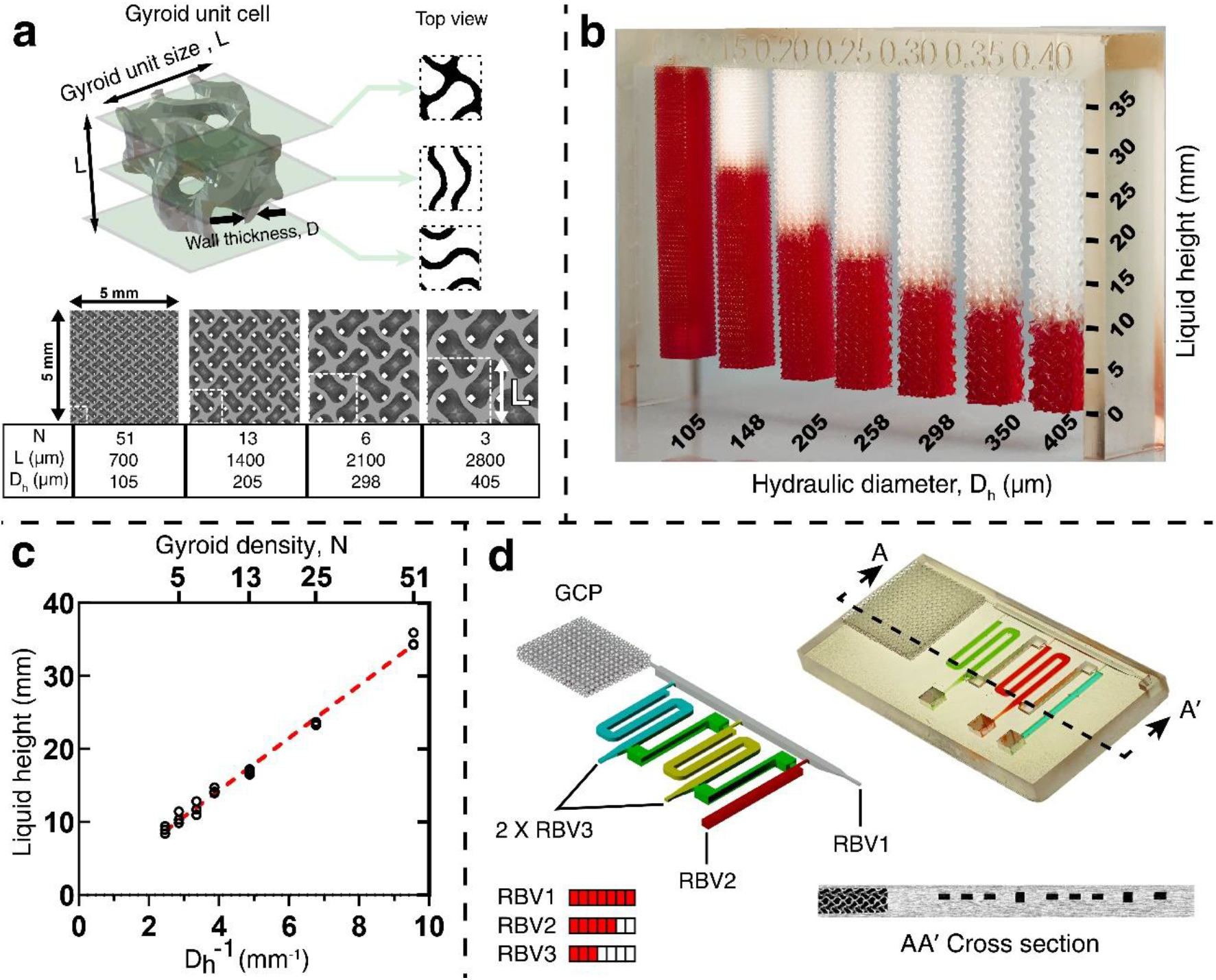
Gyroid capillary pump (GCP). **(a)** 3D image of a Gyroid unit cell and top view cross-sections at different planes. Columns of microstructures containing gyroid can be designed to achieve different capillary rise and density. **(b)** Capillary pressure and capillary rise increase with a smaller gyroid and smaller hydraulic diameter. See supplementary **Video S5. (c)** Graph of the inverse of the hydraulic diameter of the GCP versus the height of capillary rise reveals a linear relationship. **(d)** 3D-printed functional, ready-to-use CC including GCP and MCR for sequential delivery of solutions in reservoir shown in operation (**Video S6, Supporting Information**).

A Gyroid is an infinitely connected TPMS with high and regular porosity, high surface-to-volume ratio, and low flow resistance, which are advantageous for a capillary pump positioned at the outlet of a CC and designed to provide a negative pressure irrespective of the pumped volume. Gyroid units of varying sizes, ranging from 700 μm to 2800 μm can be designed using mathematical models to control the porosity and hydraulic diameter to encode the desired capillary pressure (**Figure 6a**). The hydraulic diameter (D_h_) for a gyroid structure can be determined using the equation D_h_ = 4V/P, where V represents the unit cell volume, and P is the wetted perimeter of the fluid path. This equation provides a measure of the effective cross-sectional area for fluid flow through a channel or pore in the structure, and P and V can be obtained through mathematical models. Gyroid structures provide a large volume for draining liquids due to their high porosity. Using our hydrophilic ink, we 3D printed 5×5 mm^2^ columns of microstructures with different gyroid sizes and densities to evaluate the capillary rise. As shown in **Figure 6b**, the capillary rise increased by increasing the gyroid density and reducing the hydraulic diameter (**Video S5, Supporting Information**). The correlation between the capillary rise and hydraulic diameters of gyroid columns is consistent with Jurin’s law, which states that the height of a liquid column is inversely proportional to the tube diameter (**Figure 6c**). To show the potential of gyroids as a capillary pump to develop fully functional circuits, we designed, and 3D printed a capillary device consisting of MCR and GCP and successfully achieved sequential delivery of three reagents (**Figure 6d, Videos S6-S7, Supporting Information**). The size of the GCP was 15.5 × 14 × 2.3 mm^3^, with a porosity of 62%, and a volumetric capacity of 310 μL. Whereas the capillary pressure of GCP remains comparatively weak relative to paper, which has much smaller pore size, our results show that the GCP, coupled with a capillary circuit, could be used to run the sequential delivery of 3 reagents. Future improvements in resolution will allow making GCPs with higher capillary pressure, while modulation of pore size could be used to tune pressure continuously.

## 3. Conclusions and future works

We have introduced DM of functional microfluidic systems as monolithic CCs with embedded conduits thanks to the hydrophilic CCInk made of PEGDA-250 with AA, and that can be 3D printed using common DLP 3D printers. We improved CC reliability and functionality thanks to a new SV/TV design, conduits with circular cross-sections, and embedded GCPs. The functionality of these CCs was demonstrated with a series of multilayer CCs with sequential delivery by RBV and MCRs, and an immunoassay. Functional CCs can be created by anyone with a DLP printer with a projected pixel size ≲50 μm, thus enabling distributed DM thanks to the wide availability and low cost of 3D printers, and the ready-to-use quality of CCs. We foresee that the comparative ease-of-use and low maintenance of DLP printers will facilitate DM of microfluidic as it circumvents the need of advanced skills in neither manufacturing nor microfluidics, given that designs could simply be downloaded from an online repository (e.g. www.printables.com/@JunckerLab_743461), while making it affordable with material costs < 1 US$ per chip (depending on volume) thanks to the low cost of CCInk. We hope that the advances reported here will spur DM of microfluidics and lead to a broader adoption and exploration of microfluidics and CCs in particular, and help catalyse new ideas and applications in synthesis, analysis, assays, and diagnostics and that they will be shared as ‘3D apps’ that can be downloaded from online repositories.

## 4. Material and Methods

### 4.1. CCInk Formulation and Preparation

The photocurable inks in this study were made of the monomer poly (ethylene glycol) diacrylate (PEGDA, MW250, no. 475629; Sigma Aldrich), the crosslinkers acrylic acid AA, no. 147230; Sigma Aldrich) and methacrylic acid (MA, no. 155721; Sigma Aldrich), the photoinitiator diphenyl(2,4,6-trimethylbenzoyl) phosphine oxide (TPO, no. 415952; Sigma Aldrich) (0.5 w/w) and the photoabsorber 2-isopropylthioxanthone (ITX, no. TCI0678; VWR International) (0.8 w/w). AA and MA concentrations were varied towards identifying the optimal concentrations. The CCInks were prepared by mixing the above-mentioned components using the optimal concentration for 30 min. All inks were stored in amber glass bottles after preparation.

### 4.2. 3D printing Microfluidic Devices

All capillaric devices were designed in AutoCAD (Autodesk), exported as “STL” files, and 3D-printed with two different DLP-based 3D printers with 385 nm LED light source, the Asiga MAX X27 (GV Canada Inc., Canada), and Miicraft Prime 110 (Creative CADworks, Concord, Canada) with a projected pixel size of 27 μm and 40 μm, respectively. Unless otherwise noted, all CCs reported in this work are printed with a layer thickness of 20 μm and an exposure time of 950 ms at a light intensity of 5 mW/cm^2^. Immediately after printing, to remove unpolymerized ink and clean channels, closed channels were rinsed with isopropanol (Fisher Scientific, Saint-Laurent, Quebec, Canada), and dried under a stream of pressurized nitrogen gas.

### 4.3. Contact angle measurements

2-3 μl of DI water was placed on the top surface of 3D-printed samples and imaged using a Panasonic Lumix DMC-GH3K. The side view images were imported to ImageJ ver. 1.53 (public domain software, NIH, Bethesda, MD, USA) to measure the static contact angle using the contact angle plugin. All measurements were performed on freshly UV-cured samples after cleaning and drying 3D-printed samples. To measure contact angle on plasma-treated surfaces, the above measurements were performed immediately after treating samples for 15 s at 100% power in a plasma chamber (E50 plasma chamber, Plasma Etch, Carson City, USA).

### 4.4. Penetration depth and absorbance measurements

To modulate the surface hydrophilicity properties by adjusting the contact angle, the formulation was mixed with different amounts of AA and MA ranging from 5% (w/w) up to 15% (w/w). In order to measure the effect of crosslinker on the penetration depth of light, a drop of the formulation was placed on a glass slide and exposed to different exposure times at a light intensity of 5 mW/cm^2^. After rinsing uncured ink with 70% ethanol, we measured the thickness of patterned regions using a stylus profilometer (DektakXT, Bruker Co.).

Absorbance measurements of the ink were performed by triplicate using a NanoDrop@ND-1000 (NanoDrop Technologies, Wilmington, DE, USA).

### 4.5. X-ray Photoelectron Spectroscopy (XPS)

To confirm the presence of the COOH group, the HR-XPS (Thermo-Scientific K-Alpha, Waltham, MA, USA) was conducted on 3D printed samples (10 × 10 × 2 mm^3^).

### 4.6. Toluidine blue assay to quantify functional groups

Toluidine blue assay (TBO)^[28]^ was used to quantify the density of carboxyl groups of PEGDA co-polymerized with different acrylic and methacrylic acid concentrations. Square-shaped samples (10×10×2 mm^3^) were 3D printed and washed with isopropanol (Fisher Scientific, Saint-Laurent, Quebec, Canada) before using for the assay. PEGDA samples without hydrophilic co-monomers were fabricated under the same conditions and used as controls. To carry out the TBO assay, three samples of each concentration were incubated in 2 mL of a solution containing 1 mM toluidine blue (no. 89640; Sigma Aldrich) in PBS at pH 10 at room temperature on an orbital shaker set at 150 rpm (no. 57018-754VWR; VWR International, Brisbane, CA). After 48 h, samples were rinsed frequently with PBS at pH 10 to remove unbounded dye for 1 day. The samples were then immersed in 2.5 mL of 50% vol. glacial acetic acid in Milli-Q water for 3 h. Finally, NanoDrop@ND-1000 spectrophotometer was used to measure the absorbance at 633 nm to quantify the density of TBO.

### 4.7. COMSOL Simulation

To measure the bursting pressure at SVs, COMOSL Multiphysics v.5.5 (COMSOL AB, Stockholm, Sweden) was used to simulate two-phase capillary flow within embedded channels using the level set method. A free tetrahedral mesh with “fine” size was applied for each simulation. Atmospheric pressure was set for the outlet, and the inlet static pressure was gradually varied (with 50 Pa increments) to determine the pressure of stop valve failure. The contact angle on the channel walls and the interface thickness were set 40° and 8×10-8 m, respectively. For conventional SVs, we set the contact angle of the top wall (cover tape) to 110°. Each simulation was performed with a time period of 0-0.002 s with a time step of 2.5 ×10^−5^ s.

### 4.8. Testing capillaric devices

Printed chips were pre-loaded with dyed solutions (2% food dye in Milli-Q water) and connected to filter papers (Whatman CF4, Cytiva, Marlborough, Massachusetts, USA) serving as the paper capillary pump for conducting experiments, except when embedded 3D printed capillary pumps were used. A solution with low surface tension (2% food dye and 0.1%Tween 20 in Milli-Q water) was used to evaluate edge flow and bubble trapping in conduits with either rectangular or circular conduits, presented in **Figure 4**.

### 4.9. Covid-19 assay test on nitrocellulose membranes

Nitrocellulose lateral flow membranes (vivid™ 120, no. VIV1202503R; Pall Corporation, Port Washington, USA) were cut into 5.2×12 mm^2^ rectangles using a Silhouette Portrait cutter (Silhouette, Lindon, USA). SARS-CoV-2 nucleocapsid protein was purchased from Sino Biological, Inc. (no.40588-V08B; Beijing, China) and deposited at the concentration of 1.25 mg/ml as previously described^[12,19]^ on the membranes using a programmable inkjet spotter (sciFLEXARRAYER SX, Scienion, Berlin, Germany). Before running the assay, the strips were immersed in a blocking buffer (1% BSA and 0.05% Tween 20) for one hour on an orbital shaker (no. 57018-754VWR; VWR International, Brisbane, CA) set at 150 rpm and thereafter dried overnight at 4° Celsius. The sample solutions were prepared by spiking biotinylated SARS-CoV-2 N protein rabbit monoclonal antibody (no. 40143-R004-B; Sino Biological, Inc.; Beijing, China) in buffer solution (0.1% BSA and 0.05% Tween 20 in PBS) at the concentrations of 0, 1, 10, 100, 1000 and 10000 ng/ml. Streptavidin poly-HRP (Pierce, no. 21140; ThermoFisher; Ottawa, Canada) solutions were prepared in the buffer solution with a concentration of 5 μg/mL. SIGMAFAST™ DAB tablets (no. D4293-50SET; Sigma Aldrich; Oakville, Canada) were dissolved in Milli-Q water to prepare substrate solution. The membrane strips were connected to a glass fiber conjugate pad (G041 SureWick, Millipore Sigma, Oakville, Ontario, Canada) on one end and were sandwiched between three absorbent pads (Electrophoresis and Blotting Paper, Grade 320, Ahlstrom-Munksjo Chromatography, Helsinki, Finland) serving as the capillary pump at the other end. Absorbent pads were clamped with a paper clip.

### 4.10. Image Analysis

Nitrocellulose strips were removed from the chip after the assay completion, dried at room temperature, and scanned at 600 DPI (Epson Perfection V600). The images were then imported in Image-J ver. 1.53 (public domain software, NIH, Bethesda, MD, USA) to measure the grey value at the test line and top and bottom backgrounds located 1.5 mm below and above the test line. The signal intensity for each strip was calculated by subtracting the average background (of top and bottom) from the grey value at the test line. After that, the average signal intensity of the negative controls (0 ng/ml) was subtracted from the signal intensity, plotted and fitted to generate the normalized standard curve.

### 4.11. Bursting pressure characterization

To evaluate the bursting pressure at face-centric SVs, modules with different cross-sectional areas were 3D printed. Each module consists of a face-centric SV connected to an expansion chamber. A conical inlet/outlet was designed, and 3D printed for each SV for tubing connections and to avoid bubble formation. The modules were connected to a pressure pump (MFCS-4C) from both ends and Fluiwell package (Fluigent) with reservoirs containing 2% food dye in Milli-Q water. Once the SV was capillary-filled, MAESFLO v.3.3.1 software (Fluigent) was used to control positive or negative pressure to calculate the burst pressures of the SV with increments of 10 Pa.

### 4.12. Videos and image stacking

Videos and images of 3D-printed chips were taken using a Panasonic Lumix DMC-GH3K and Sony α7R III. For focus stacking, Imaging Edge Desktop (Sony Imaging Products & Solutions Inc., Japan) was used to take the sequence of images on different focal planes. Then, CombineZP (available at https://combinezp.software.informer.com/) was used to process the images. Micro-computed tomography (Micro-CT) was performed using Skyscan 1172 (Bruker, Kontich, Belgium) at a pixel size of 5 μm, and used to confirm the dimensions of embedded channels.

## 5. Acknowledgements

We would like to thank Bishakh Rout for conducting XPS measurements presented in the Supporting Information. A.S.K. acknowledges postdoctoral fellowship no. 267919 from the Fonds de recherche du Québec – Nature et technologies (FRQNT). V.K. acknowledges FRQNT for the doctoral scholarship (no. 268838). D.J. acknowledges support from a Canada Research Chair in Bioengineering.

## 6. Contributions

V.K. and A.S.K. contributed equally to the project. V.K. formulated the materials, developed the Gyroid capillary pump concept, and made contributions to the original draft, while A.S.K. designed the CCs, developed face-centric stop valves, conducted simulations, and prepared the initial draft. Both V.K. and A.S.K. contributed equally to the remainder of the project by performing experiments, analysing data, characterizing formulations and optimization, conducting on-chip assays, capturing images and videos, and reviewing and editing the manuscript. M.S. assisted with the assay by spotting antibodies on the nitrocellulose membranes. D.J. provided funding, supervised the project, and contributed to writing the original draft, reviewing, and editing the manuscript.

