## Supplementary Information for "Digital Manufacturing of Functional Ready-to-Use Microfluidic Systems"

1. Water contact angle on plasma-treated 3D-printed samples over time
2. Absorption of photocurable ink
3. Previous fabrication method for capillary microfluidic devices
4. XPS results

### Supplementary Videos

**Video S1.** Multilayer Microfluidic Chain Reaction (MCR)

**Video S2.** CC with circular channels

**Video S3.** Micro-CT of CC with circular channels

**Video S4.** Autonomous assay using a 3D-printed monolithic CC

**Video S5.** Capillary rise in gyroid columns

**Video S6.** CC with GCP and MCR

**Video S7.** Micro-CT of CC with GCP and MCR

**Supplementary Figures**

**Figure S1**

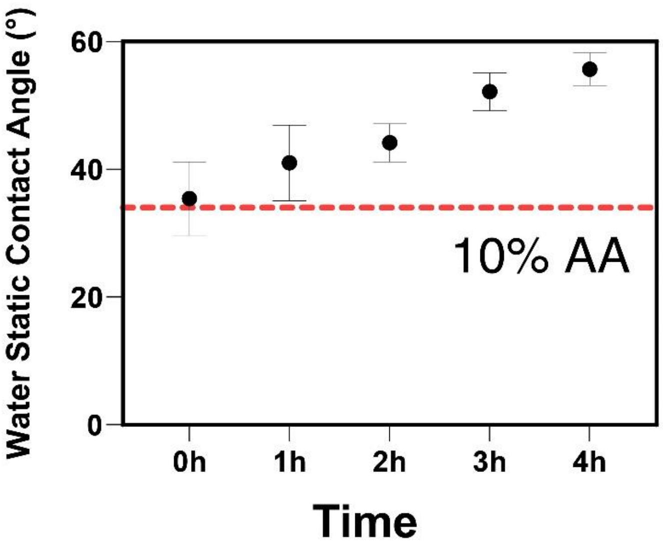


**Figure S1. Water static contact angle on plasma-treated 3D printed samples over time.** The contact angle changes by approximately 20$^o$ over a period of 4 h.

**Figure S2**

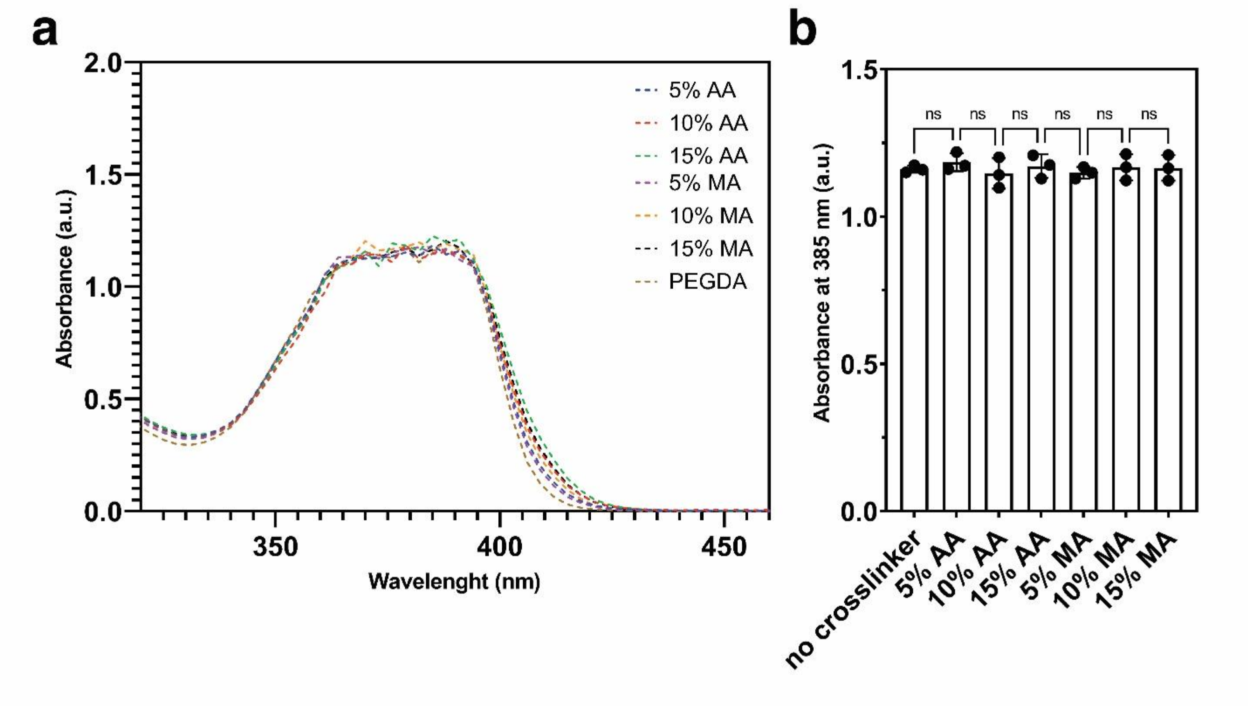


**Figure S2. Impact of additives on ink absorption at 385 nm illumination wavelength of DLP projector. (a)** Absorption of CCInk at 385 nm for different CCInks with varying additive concentrations. **(b)** Absorption of CCInk at 385 nm. Crosslinker concentration does not affect the light absorption of the ink. This is due to low absorption of AA and MA compared to PI and PA.

**Figure S3**

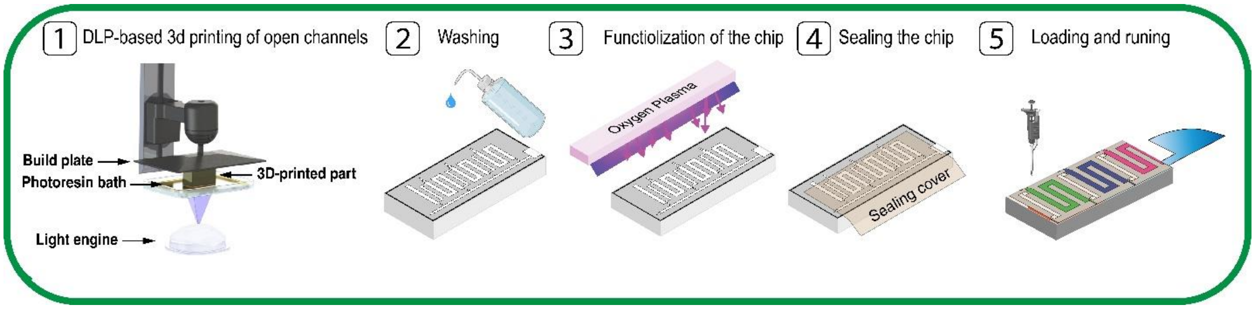


**Figure S3. Previous fabrication method for capillary microfluidic devices.** 1) 3D printing of open channels. 2) Removal of uncured ink. 3) Plasma-treating of the chip. Hydrophilization of CCs using a plasma chamber results in transient hydrophilicity, lacks reproducibility, and limits the design of CCs to open surface channels that are sealed with tape. 4) Sealing the chip with a hydrophobic transparent tape. 5) Loading the chip.

### Figure S4

Without acrylic acid (98.7% PEGDA+ 0.5% TPO + 0.8%ITX)

| Name | Peak BE | FWHM eV | Area (P) CPS.eV | Atomic % |
|---|---|---|---|---|
| O-C=O (Carboxylic acid) | 288.43 | 0.57 | 130.23 | 1.4 |
| C-O-C | 285.97 | 1.06 | 1902.9 | 20.45 |
| C-C | 284.36 | 1.4 | 7280.16 | 78.15 |

With acrylic acid (98.7% PEGDA+ 0.5% TPO + 0.8%ITX+ 5% AA)

| Name | Peak BE | FWHM eV | Area (P) CPS.eV | Atomic % |
|---|---|---|---|---|
| O-C=O (Carboxylic acid) | **288.36** | **0.84** | **522.56** | **5.09** |
| C-O-C | 285.96 | 1.22 | 2462.6 | 23.93 |
| C-C | 284.4 | 1.33 | 7311.81 | 70.98 |

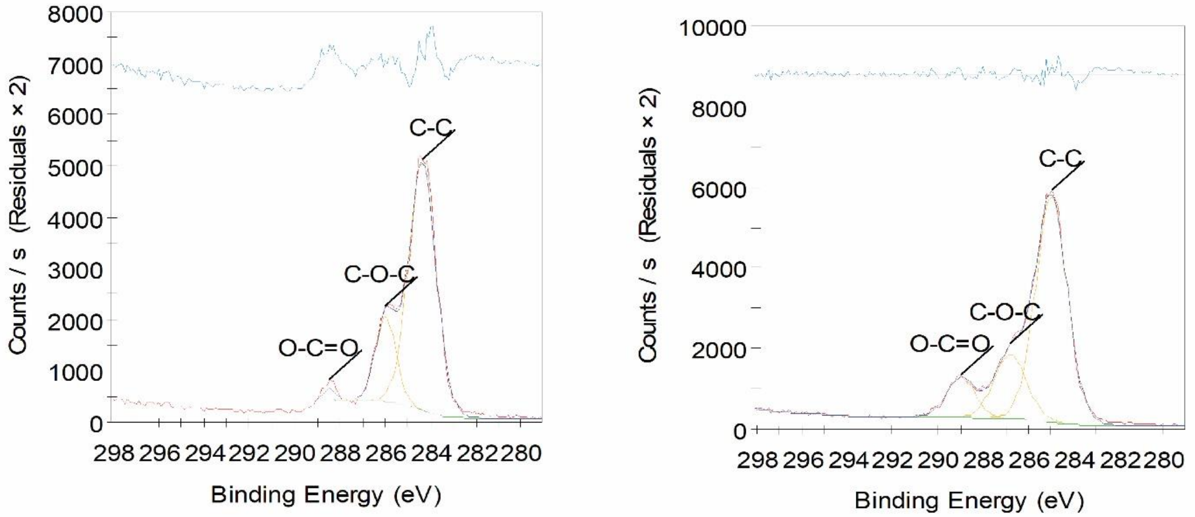


**Figure S4. X-ray photoelectron spectroscopy (XPS) measurement on the samples with and without AA**. The results showed a presence of approximately 5.1% carboxylic acid (COOH) group in the samples with 5% AA, while samples without AA only exhibited approximately 1.4% COOH.
